# Boundary Vector Cells Encode a Future-Biased Spectrum of Positions in the Rat

**DOI:** 10.64898/2026.01.11.698891

**Authors:** Ehren Lee Newman, Inna Mashanova-Galikova, Zoran Tiganj, Colin Lever

## Abstract

Spatial tuning is a hallmark property of neural firing in the hippocampal formation. Yet, that tuning is often less well correlated with the instantaneous current position of an animal than it is with an integrated version of the past or future state of the animal. Whether that encoding is biased towards past or future states and the extent to which it shows fixed versus multi-scale encoding varies across circuits and cell types. The temporal encoding properties of boundary vector cells of the subiculum are not well established. To address this here, we re-analyzed recordings of BVCs described previously by Lever et al. (2009) with multiple approaches. In the first, we asked if adding a temporal offset between the rat position and the spiking of a BVC increased the apparent spatial tuning in the firing rate map. We found that aligning BVC spiking with future states maximized the rate map spatial tuning. These results were mirrored in a second analysis that, instead of optimizing rate map spatial tuning, optimized how well the firing rate map predicted the BVC spiking. The second analysis also allowed us to ask whether that encoding is focused on a particular temporal horizon or whether the encoding captures behavior at multiple scales. To this end, for a given recording, we asked “How much time-integration of the behavioral state is the observed spiking most consistent with?” We observed a wide spectrum of time-constants of integration across cells, indicating that BVCs form a multiscale encoding of future states. The distribution of both offsets and integration rates observed across BVCs did not differ significantly from other, non-BVC, subiculum neurons. Taken together, these findings indicate that BVCs, along with other subiculum neurons, form a multi-scale encoding of future states.

## 1 Introduction

Cellular and oscillatory activity in the hippocampal formation exhibits robust correlations with navigational states, including position (O’Keefe and Dostrovsky, 1971; Ekstrom et al., 2003; Ulanovsky and Moss, 2007; Hafting et al., 2005), head direction (Taube et al., 1990), boundary proximity/direction (Lever et al., 2009; Solstad et al., 2008; Alexander et al., 2020), and locomotion, including running speed (Sargolini et al., 2006; Caplan et al., 2003; Wells et al., 2013). These correlations are typically derived by comparing the neural variable in question to the subject’s instantaneous state, such as the firing rate of a place cell to the instantaneous position of the rat locomoting in an open field. Intriguingly, however, such neural correlations are often enhanced when spiking activity, for example, is aligned not with the animal’s instantaneous state, but rather with its time-shifted or time-averaged state (Muller and Kubie, 1989; Blair and Sharp, 1995; Dannenberg et al., 2019; Bright et al., 2020; Chaudhuri-Vayalambrone et al., 2023).

For example, the spatial tuning of CA1 neurons is more precise if aligned with the animal’s position around 120 ms after the spike occurred, instead of its position at the time of the spike (Muller and Kubie, 1989; Chaudhuri-Vayalambrone et al., 2023). Similarly, spatial tuning in the subiculum (Sharp, 1999) and in grid cells of the medial entorhinal cortex (Chaudhuri-Vayalambrone et al., 2023) is enhanced by aligning spikes to future positions. These observations are consistent with the idea that the entorhinal-hippocampal system is generating predictions about future states (Stachenfeld et al., 2017; Momennejad, 2020; Geerts et al., 2023).

Interestingly, however, not all coding in the entorhinal-hippocampal system appears to be prospective. For instance, entorhinal speed cells, though obviously linked to locomotion, seem to provide a retrospective record, encoding the historical running speed of the animal (Dannenberg et al., 2019). Furthermore, cells in the lateral entorhinal cortex also track the recent past, coding elapsed time since salient event boundaries (Bright et al., 2020; Tsao et al., 2018). Accordingly, it is not straightforward to suggest that entorhinal-hippocampal codes are simply predictive.

Beyond the prospective-retrospective dimension, a further important dimension of functional interest concerns multiple timescales. For instance, considering prospective coding, entorhinal grid cells encode the rat’s future position with temporal offsets proportional to their spatial scale, thereby providing a multiscale representation of the animal’s trajectory (Chaudhuri-Vayalambrone et al., 2023). Considering retrospective coding: speed cells encode the historical running speed of the animal at widely-differing timescales ranging from hundreds of milliseconds to hundreds of seconds, depending on the neuron (Dannenberg et al., 2019); and lateral entorhinal cells encoding the amount of time that has passed since critical event boundaries at multiple timescales (Bright et al., 2020; Tsao et al., 2018).

Theoretical modeling shows that multiscale encoding provides a rich basis upon which to form memories and predict future states (Howard et al., 2014; Shankar et al., 2016). A key prediction of that modeling is the presence of a spectrum of encoded timescales. Such a spectrum enables encoding of a memory timeline of *what happened when* and forming an estimate of the timeline of the future. The multiscale encodings observed in entorhinal neuronal activity (e.g., Chaudhuri-Vayalambrone et al. 2023; Bright et al. 2020; Tsao et al. 2018; Dannenberg et al. 2019) exemplify functional types of spectral encoding. Importantly, in this theory, the type of information encoded determines the types of inferences obtainable. For example, multiscale encoding of time since event boundaries supports retrospective reconstruction of a timeline.

With these considerations of both the prospective-retrospective and the multiscale dimensions in mind, here we examined temporal coding in the boundary vector cells (BVCs) of the subiculum (Lever et al., 2009). The existence of such cells was first predicted by computational models of inputs to place cells (O’Keefe and Burgess, 1996; Hartley et al., 2000; Lever et al., 2002a) seeking to explain how place cell firing fields are shaped by environmental geometry (O’Keefe and Burgess, 1996; Lever et al., 2002b). BVCs fire when an environmental boundary (such as a wall or drop edge) or sufficiently large object, is at a specific distance and allocentric direction from the animal (Lever et al., 2009; Stewart et al., 2014; Poulter et al., 2021). BVCs show some plasticity: for example, their firing fields can be influenced by the (e.g., rectangular) geometry of the environment (Muessig et al., 2024); a subtype of BVCs, called vector trace cells, also exhibit *memory* for the locations of boundaries and objects (Poulter et al., 2021). Such rich encodings

Generally, less is known about subicular BVCs than about other cell types in the entorhinal-hippocampal system, see (Poulter et al., 2018; Bicanski and Burgess, 2018, 2020; Lever et al., 2025) for reviews and discussion. Whether subicular BVCs encode boundary vectors retrospectively or prospectively, and whether this code operates over a range of timescales, are open questions. To address these questions, we first applied rate-map methods analogous to those used to reveal multiscale temporal coding in grid cells (Chaudhuri-Vayalambrone et al., 2023). We then used maximum-likelihood estimation to determine, for each cell, whether spikes were better predicted by time-shifted and/or time-integrated representations of the rat’s position. The results demonstrate that spatial tuning for typical BVC firing aligns most strongly with time-averaged estimates of *future* position but that, across cells, there is a spectrum of offsets and temporal scales encoded.

## 2 Methods

New analyses were performed on recordings obtained by and first described by Lever et al. (2009). Subjects, surgery, and data collection are paraphrased here for ease of reference.

### 2.1 Subjects and Surgery

Data were obtained from six male Lister Hooded rats, weighing 315-390g at time of surgery. Rats were maintained on a 12:12 hour light:dark schedule (with lights off at 15:00). Food deprivation was maintained such that subjects weighed 85-90% of free feeding weight. Recordings were made from the dorsal subiculum. Details of the implantation and confirmation of electrode placement were described previously (Cacucci et al., 2004; Lever et al., 2009). Briefly, a bundle of four HM-L coated platinum-iridium wire tetrodes mounted to a microdrive was chronically implanted dorsal to the subiculum under deep anesthesia. Tetrodes were lowered to isolate cells after surgery.

### 2.2 Electrophysiology, Location and Head Orientation

Tetrodes were lowered towards the subiculum region and then left to stabilize before the start of the recording. The head stage was connected to a pre-amplifier by 3-meter lightweight wire. The outputs of the pre-amplifier passed through a switching matrix, and then to the filters and amplifiers of the recording system. Each channel was continuously monitored at a sampling rate of 50 kHz and action potentials were stored as 50 points per channel whenever the signal from any of the 4 channels of a tetrode exceeded a given threshold. EEG signals were amplified 10-20K, band-pass filtered at 0.34-125 Hz and sampled at 250 Hz. (See details in Lever et al. 2009).

Head position and orientation were tracked using an overhead video camera and tracking hardware/software by tracking the position of two arrays of head-mounted small, infrared LEDs, one array brighter and more widely projecting than the other. Cluster cutting to isolate single units was performed manually using custom made software (TINT, Axona, UK). (See details in Lever et al. 2009).

### 2.3 Experimental Design and Training

Recordings were collected as rats foraged for cereal in multiple distinct open field arenas. Rats were transferred to the arenas in a consistent fashion to maintain orientation. At the end of each trial, the rat was removed from the recording environment, and placed back on the holding platform until the next trial. The length of the trial was proportional to the size of the arena and was in the range 10-16 minutes. The inter-trial interval was 20 minutes. The arena specifics are described by Lever et al. (2009). Briefly, they varied with regard to shape (square in Envs. ‘a’ & ‘d’ and circle in Envs. ‘b’ & ‘c’), size (‘a’: 62 cm; ‘b’: 79 cm; ‘c’: 90 cm, ‘d’: 39 cm across), and boundary type (morph-box walls in Env. ‘a’, smooth in Env. ‘b’ & ‘d’, and the platform edge in Env. ‘c’). A curtain surrounded the testing arena in all conditions but Env. ‘b’. For some cells, additional recordings were performed in the dark in Env. ‘a’, with the box rotated 45^◦^, or with two morph-boxes combined and a 60 cm long barrier was placed halfway along the north-south or east-west midline. In all recordings, an external white cue card (102 cm high, 77 cm wide) provided directional constancy.

### 2.4 Data Inclusion and Exclusion Criteria

The BVC dataset here was comprised of 194 unique recordings, representing an average of six trials for each of 33 unique BVC cells characterised in the original report of BVCs (Lever et al., 2009). Where indicated, cells from tetrodes located in the subiculum but not labeled as BVCs were also analyzed (i.e., non-BVCs). The non-BVC dataset was comprised of 377 unique recordings, representing an average of about five trials for each of 77 unique cells.

### 2.5 Ratemap Creation

Firing ratemaps were constructed from 2.6 x 2.6 cm binned data, smoothed using a Gaussian smoothing kernel with a standard deviation of 5.2 cm. Spike count was divided by occupancy time to obtain a firing rate per bin. To display firing ratemaps in figures, the ratemaps were auto-scaled false color maps, each color representing a 15% band of peak firing rate, from dark blue (0-15%) to red (85-100%). Peak rate (after smoothing) is shown next to each ratemap.

### 2.6 Ratemap Tuning Based Approaches Estimating Time-Shift

Following previously used approaches for estimating the temporal lag of spatial coding, we examined results from zero-lag spatial autocorrelation (ZLAC), (Chaudhuri-Vayalambrone et al., 2023) and spatial information (s.i.) (Skaggs et al., 1993) approaches with variable temporal offsets between the tracking and spiking. In both approaches, we analyzed each recording for each cell separately. In doing so, we shifted the timing of the spiking relative to the tracking by two seconds into the past and into the future one video frame at a time (i.e., 20 ms steps given the 50 Hz video sampling). For each step, we created a ratemap and then computed the ZLAC and s.i. scores yielding a curve of scores over offsets. We then identified the ‘peak shift’ as the local maxima closest to zero shift after smoothing with a boxcar kernel with a width of 6 (120 ms). This was done separately for ZLAC and s.i.. Significance of ZLAC and s.i. at any lag was established by performing 500 pseudo-random rotations between the tracking and spiking. The pseudo-random rotations were constrained to shift the relative alignment of the data types by at least 5% of the recording length (30 sec in the case of the 10 min recordings). The ‘rotation’ here is the wrapping of the misaligned data from one end of the recording around to the other (e.g., if the behavior was shifted earlier by 120 s, the 120 s that now ‘occurs’ before the electrophysiology recording was appended to the end of the behavioral tracking that would otherwise end before the end of the electrophysiology). The permuted ZLAC and s.i. scores were used to compute a z-score type normalization of the empirical ZLAC and s.i. values, respectively. Normalized values were considered to reflect significant spatial tuning if they were larger than 3.4808, reflecting *p <* 0.00025 (the Bonferroni corrected *p <* 0.05 over 200 comparisons). In this work, we used spatial information computed as a function of bits per spike. Note, however, that because the optimization was performed for each recording separately, preserving the recording duration and total spikes, it would not have changed the qualitative results had it been performed with the bits per second variant.

### 2.7 Maximum Likelihood Estimation of Spatio-Temporal Tuning

The ratemap tuning based approach described above maximizes the amount of spatial tuning in a ratemap without consideration for whether that ratemap accurately captures the firing statistics of the neuron. Addressing this, we also applied an analysis approach that sought to identify parameters that accurately capture the firing statistics. For this, we used a maximum likelihood based approach: Temporal tuning parameters were selected so as to maximize how well the resulting firing ratemap could be used to predict the moment by moment spiking of the neuron.

In this approach, the quality of a given set of parameters was assessed as follows:

1. Compute a surrogate of the time-varying position of the rat as

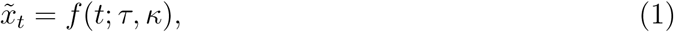 where *τ* is the magnitude and direction (future or past) of a time-shift and *κ* controls the width of a time integration window. How this was done specifically differed over analyses and is unpacked for each specific analysis below.
2. For a given set of surrogate *x_t_*values, build a ratemap that per the methods described above.
3. Compute the probability of observing a spike in each time-bin by using the ratemap as a look-up table of the expected firing rate based on the corresponding *x_t_* value and converting the firing rates into the probability of observing a spike using the Poisson probability mass function,

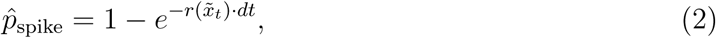 where *r*(*x̃*) is the ratemap value at surrogate position *x̃* and *dt* is the bin-width used in discretizing the predictions, set to 0.02 sec (one frame of the tracking).
4. Compute the total negative log-likelihood (NLL) for the entire time series. Because the model treats each time bin as an independent Bernoulli trial, the total NLL is the sum of the NLLs for each individual bin. This is calculated as:

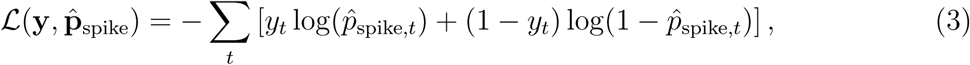

where the sum is over all time bins *t* in the recording, **y** is the vector of observed binary spike data (where *y_t_* ∈ {0, 1}), and **p̂**_spike_ is the corresponding vector of the model’s predicted spike probabilities. This calculation yields a single scalar value that quantifies the goodness-of-fit for a given set of parameters, *τ* and *κ*. Lower L(**y**, **p̂**_spike_) values indicate better predictions of the observed spike train.

This approach enables testing of whether a given parameter significantly improves how well the ratemap obtained with given values of *τ* and *κ* predicts the observed spiking. Broadly, this is done by testing whether L(**y**, **p̂**_spike_) is significantly better when a parameter is treated as a free-parameter (i.e., one that can be optimized via a gradient descent algorithm) instead of being fixed to a default value. Details of statistical testing are described below.

#### 2.7.1 Time-Shift

The time-shift parameter, *τ* , temporarily offsets the spiking with respect to the behavioral tracking. A value of 0 corresponds to how standard ratemaps, with no temporal shift, are constructed. Positive *τ* values align spiking with future positions (i.e., predict spiking from where the rat will be). Negative values align spiking to past positions (predict spiking from where that rat was). We tested whether optimizing *τ* generated ratemaps that reliably predicted spiking better than those generated with no time-shift, and if so, whether that offset was toward the future or the past on average as described in the section Statistical Testing.

#### 2.7.2 Time-Integration

Time-integration estimates each position based on a time-integrated version of position over a recent window with width proportional to *κ*. The form of the time-integration depends on the shape of the kernel used. We compared three kernel shapes: Gaussian, boxcar and exponential. Each was compared to a baseline model with no integration.

The generalized form for the smoothed estimate at time *t* is:

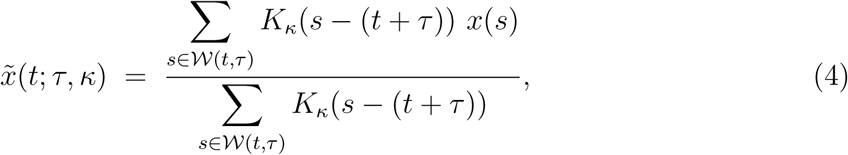

where *K_κ_*(*x*) is a given kernel function with width controlled the parameter *κ*. To prevent information leakage between training and test folds during cross-validation, the integration window W(*t, τ* ) was bounded to include only samples within 15 seconds of the shifted time point *t* + *τ* : where Δ*t* is the time bin width equal to one video tracking frame (20 ms). For all models, if a kernel’s effective window extended beyond this bound, it was truncated to W(*t, τ* ) and renormalized by the denominator in Eq. 4.

##### Gaussian (“Normally weighted”) moving average

This model tests the hypothesis that neural activity is maximally sensitive to behavior at latency *τ* , with decreasing sensitivity for times further from this point. This is implemented with a Gaussian kernel:

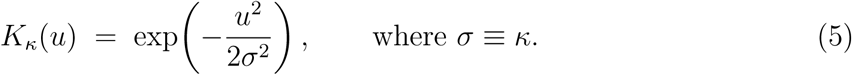

This symmetric kernel integrates states both before and after the central time point *t* + *τ* . Such a formulation is consistent with theoretical work on scale-invariant representations of time that can be adapted to encode space O’Keefe and Burgess (1996). During optimization, the standard deviation *σ* was bounded to [0.01, 100] s.

##### Boxcar (uniform) moving average

This model implements the simpler hypothesis that a neuron is equally sensitive to all behavioral states within a defined window. This corresponds to a uniform kernel applied retrospectively from the time point *t* + *τ* :

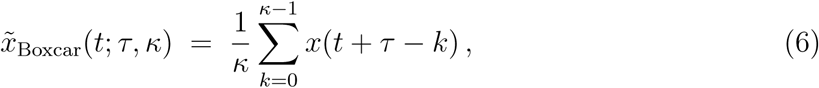

where the window width, *κ*, is the number of time bins included in the average. During optimization, *κ* was bounded to 15000 ms.

##### Exponentially weighted moving average

To capture potentially asymmetric temporal sensitivity (i.e., a retrospective or prospective bias), we used a one-sided exponentially decaying kernel. The model’s direction (past vs. future) and decay rate were governed by a single fitted parameter, *ω*, which was bounded to [−12, 12]. This parameterization avoids the numerical instability that can arise when fitting the direction and decay rate separately. Specifically, we mapped *ω* to a decay constant *κ* and a direction *d*:

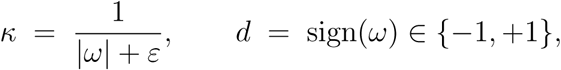

where a small constant *ε* ensures a minimal decay rate. The estimated behavioral state is then given by:

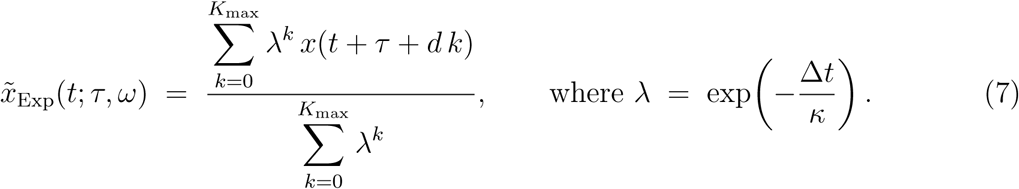

Here, *d* = +1 applies the kernel to future samples (prospective model), while *d* = −1 applies it to past samples (retrospective model). The summation is truncated at 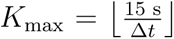 to enforce the 15000 ms bound.

#### 2.7.3 Parameter Optimization

To optimize the free parameters for each model, we used a particle swarm gradient descent approach. Specifically, we used the Matlab function particleswarm.m with SwarmSize set to 50 and HybridFcn set to fmincon, which implements a local gradient-based search after the initial swarm exploration. To ensure robust fitting, a random restart procedure was used wherein the particle swarm was reinitialized iteratively until 5 consecutive restarts failed to produce an improved fit. The parameter set that yielded the lowest final negative log-likelihood (L(**y**, **p̂**_spike_)) on the training data was kept for subsequent analysis.

### 2.8 Cross-validation

To test whether optimized parameters were generalizable to data not used in their estimation, and thereby support the critical evaluation of the goodness of the model fits, we used a K-fold cross-validation approach with *K* = 10. That is, recordings were split into 10 consecutive segments along the time variable. On each of ten separate runs, a different segment was omitted from the fitting process and the remaining *K* − 1 data segments were used for parameter estimation. The omitted segment was used as the validation dataset, used to assess the quality of the resulting fit.

Time-shifting and time-integrating procedures can blur the boundaries between the fitting and validation segments and introduce segments at the start or end of the recording without aligned behavior and spiking. To address this, buffers were cut from the start and end of the recording and around the validation dataset. Critically, a buffer of fixed size was used across all parameterizations. This was so that the number of omitted data points was constant across parameterizations and the likelihood score could not be affected by varying numbers of samples across parameter values.

### 2.9 Statistical Testing

We employed different statistical approaches across analyses as follows: We used the non-parametric Wilcoxon signed-rank test to test for significant improvements in the ratemap–based analyses. For the maximum-likelihood (ML) models we used the Bayesian Information Criterion (BIC) for within-dataset model selection. We assessed population-level effects with a General Linear Mixed-Effects (GLME) model. We assessed parameter estimate reliability with intra-class correlations (ICC). Each are described here.

#### 2.9.1 Ratemap–based analysis

We used the Wilcoxon signed-rank test to determine whether the median temporal lag that maximized spatial information (s.i.) or zero-lag autocorrelation (ZLAC) differed from zero across recordings.

#### 2.9.2 Maximum-likelihood model selection

For each recording, we compared candidate ML models using the Bayesian Information Criterion (BIC), computed from the *training-set* likelihood:

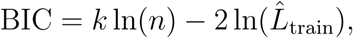

where *k* is the number of free parameters, *n* is the number of time bins, and *L̂*_train_ is the maximized likelihood on the training data. In our design, the base model had no free parameters, models with either a time shift (*τ* ) or a time-integration window (*κ*) added one free parameter, and models with both added two. We considered ΔBIC *>* 10 as strong evidence favoring the model with the lower BIC.

#### 2.9.3 Population-level inference with GLME

To ask whether particular temporal components (time shift and/or the choice of integration kernel) reliably improved predictive performance across a cell population (BVCs or non-BVCs), we fit a General Linear Mixed-Effects (GLME) model to the *cross-validated test-set* negative log-likelihood (NLL), reducing sensitivity to overfitting:

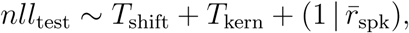

where *T*_shift_ ∈ {0, 1} indicates whether a free time-shift parameter was included, and *T*_kern_ ∈ {none, boxcar, exponential, normal} indexes the temporal-integration kernel. The model included a random intercept for the mean firing rate of the cell, *r̄*_spk_, to allow baseline differences in NLL to vary with overall firing rate. This mixed-effects model provided a superior fit compared to a fixed-effects-only model (BIC = 1.02×10^5^ vs. 1.48×10^5^), although the qualitative pattern of results did not depend on including the random effect.

#### 2.9.4 Parameter Reliability with ICC

To establish whether parameter fits were reliable (i.e., consistent) across cross-validation folds and recordings of the same cell, we used intra-class correlation (ICC) (Shrout and Fleiss, 1979; Koo and Li, 2016). We used the form to test for one-way random effects, absolute agreement, multiple raters/measurements (i.e., ICC(1,k) approach). To test for stability at the broadest level, all parameter estimates for a given cell (10 k-fold fits for each of the separate recordings of the same cell) were treated as independent raters. To test for stability across recordings of the same cell, only one of the K-fold estimates were used from each of the recordings (the first) for each cell. Per convention, values less than 0.5 indicate poor reliability, between 0.5 and 0.75 indicate moderate reliability, between 0.75 and 0.9 indicate good reliability, and greater than 0.90 are indicative of excellent reliability (Koo and Li, 2016).

## 3 Results

### 3.1 BVC spatial tuning is greater for future positions

We examined how the quality of the spatial tuning ratemap changed as a function of how far we shifted the behavioral tracking from the recorded spiking. Following approaches used previously, we identified the temporal shift that yielded a ratemap with maximum spatial information (s.i.; Sharp 1999) and the shift that yielded the maximum zero-lag auto-correlation (ZLAC; Chaudhuri-Vayalambrone et al. 2023). Figure 1 illustrates this process for five representative BVCs (Figure 1A,B). For each cell, ratemaps were created for shifts ranging between -2000 ms and +2000 ms in 20 ms intervals (only a subset of which are shown in Fig 1) and, for each, we computed s.i. and ZLAC. The resulting s.i. and ZLAC scores were then z-scored based on values obtained from 500 random shifts of 30 s or longer. Finally, we identified the temporal lag at which each measure reached its maximum. Equivalent timeshift results for simultaneously-recorded non-BVCs are shown in Figure 1C,D.

**Figure 1:**
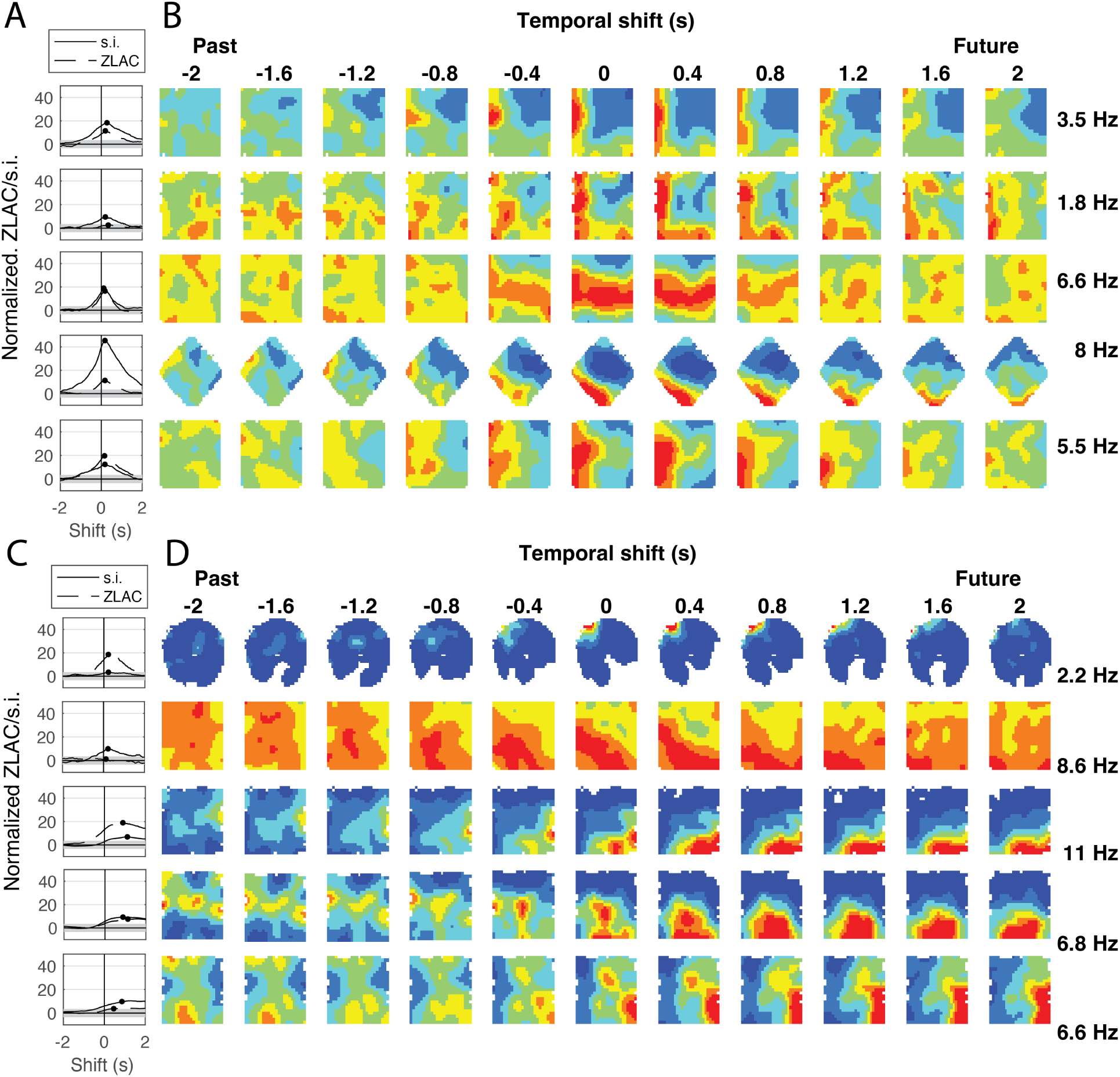
Time shifting spikes relative to rat position improves spatial tuning. A) Normalized (z-scored) spatial information (s.i.; solid line) and normalized zero-lag auto-correlation(ZLAC; dashed line) for five representative BVCs are shown plotted as a function of the temporal shift between the spiking and spatial position. B) Spatial firing ratemaps for a subset of the points shown for each of the aligned curves in A. A fixed color map is used across all ratemaps scaled to the peak firing rate observed at any shift, indicated at the right of each row. B-C) same as A-B but for five representative non-BVCs recorded in the subiculum.

Our first pass version of this analysis identified the lags that generated the maximum s.i. and ZLAC for each of the 194 individual recordings from 33 BVCs from Lever et al. (2009)) that had z-scored s.i. or ZLAC scores of at least 3.4808, corresponding to a significance threshold of *p <* 0.05 with Bonferroni correction for the 200 tests performed across lags for each cell. The temporal lags that generated the maximum z-scored s.i. were reliably positive across the 167 sessions with significant spatial information (Median = 120 ms, *W* = 12331*, z* = 10.3*, p <* 1 × 10^−24^). Likewise for ZLAC, temporal lags that generated the maximum values were reliably positive across the 171 sessions with significant ZLAC scores (Median = 140 ms, *W* = 13186*, z* = 10.3*, p <* 1 × 10^−24^). These results are shown in Figure 2. The observation that the temporal lags that result in ratemaps with maximum spatial tuning are reliably positive indicates that BVCs code for future positions.

**Figure 2:**
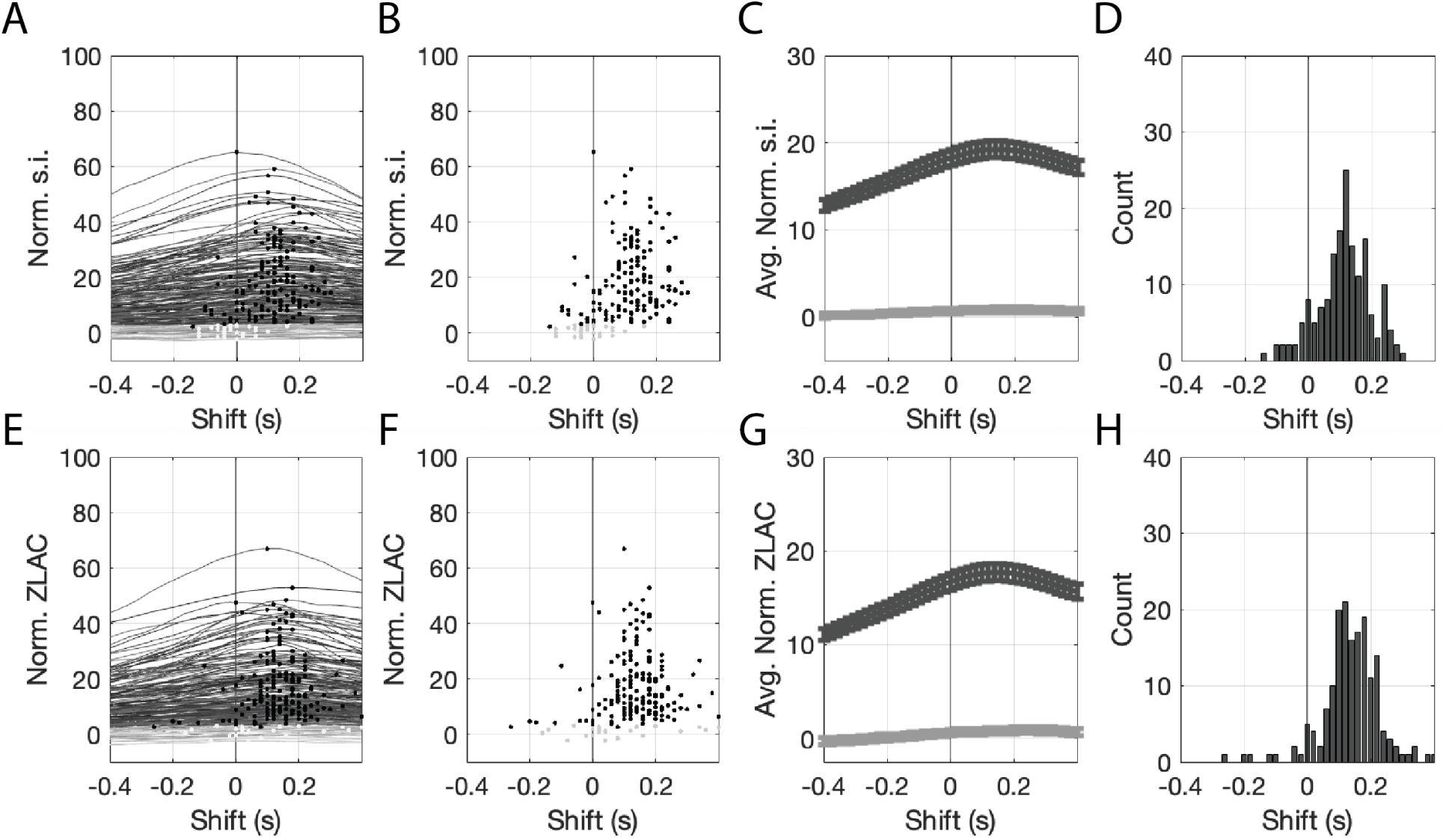
Temporal lags that generate ratemaps with maximum normalized spatial information (s.i.) and zero-lag autocorrelation (ZLAC), are reliably positive. A) Normalized s.i. plotted as a function of lag separately for each recording of each BVC. Curves with at least one significant s.i. score are plotted in black, the remainder are gray. The peak of each curve is marked by a dot. B) Dots marking the maximum normalized s.i. values, replotted from A for clarity. C) The means of the significant (black) and non-significant (gray) curves from A are shown with standard error confidence intervals. D) Histogram of the lags that resulted in the maximum normalized s.i. scores. E-H) Same as A-D but for ZLAC instead of s.i..

**Figure 3:**
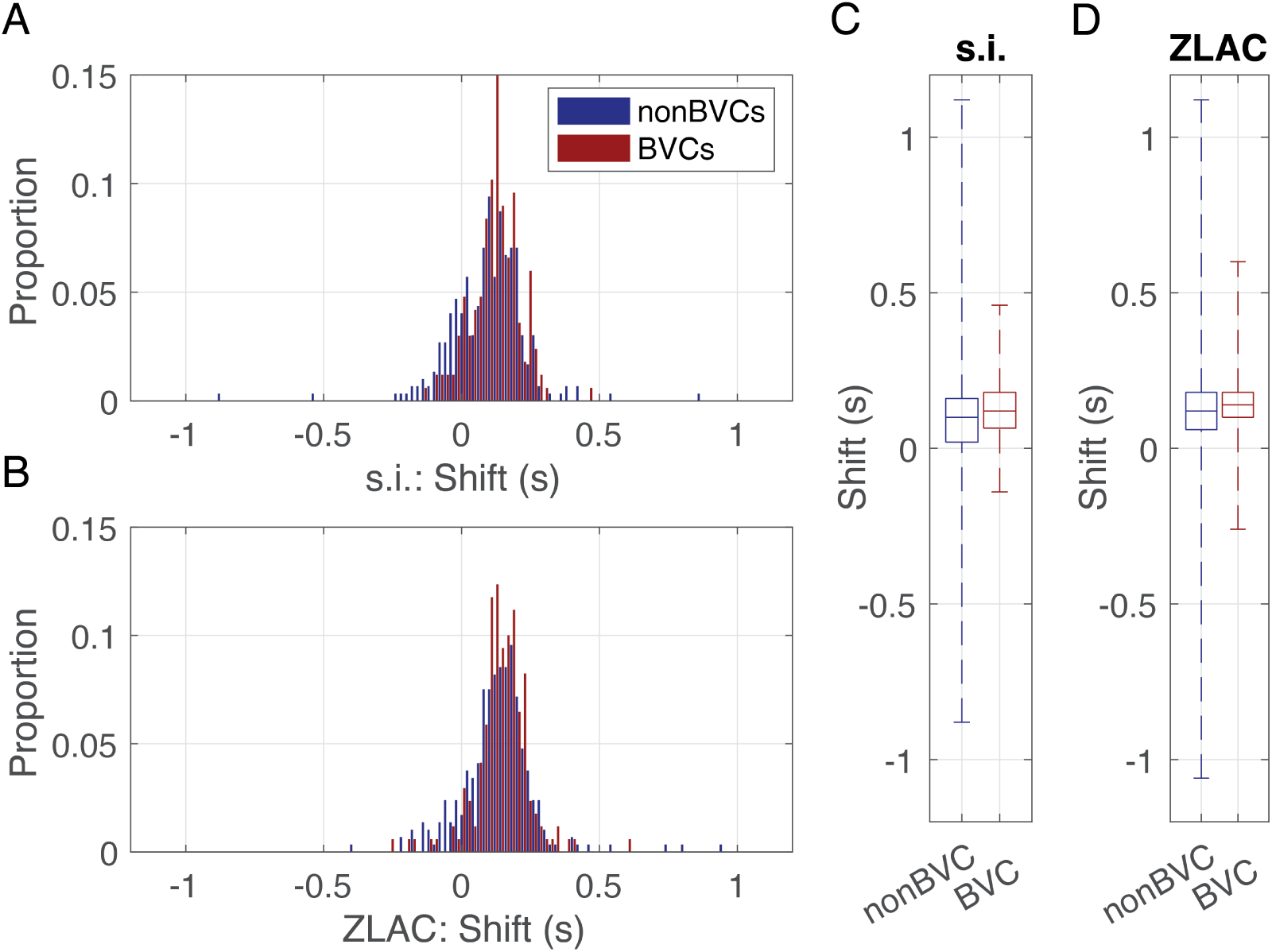
BVCs and non-BVC ratemap spatial tuning was maximized as similar temporal shifts. A) Normalized histograms of shifts that yielded the maximum ratemap spatial information (s.i.) are shown for non-BVCs (blue) and BVCs (red). B) Same as A but for shifts that yielded the maximum zero-lab auto-correlation (ZLAC). C) Box-and-whisker plot comparisons of non-BVCs BVCs in terms of the shifts that yielded the peak s.i. score. No significant difference was found. D) Same as C, but for shifts that yielded the peak ZLAC score. No significant difference was found. spiking of a cell given knowledge of the behavior? To address this, we used a maximum likelihood based approach wherein temporal tuning parameters were selected based on how well they enabled us to predict the observed spiking based on a ratemap constructed with those parameters.

To gain perspective on the temporal tuning of these BVCs, we examined the tunings of other cells coincidentally recorded on tetrodes that had BVCs. We refer to these as the non-BVCs. There were 77 unique non-BVCs, each recorded about 4.9 times each, yielding 377 recordings. Of these recordings, 298 (79%) generated ratemaps with significantly more spatial information than chance, accounting for 73 of the 77 non-BVCs (94.8%). Similarly, the ratemaps of 293 (78%) had significantly above chance ZLAC scores, accounting for 71 of the 77 non-BVCs (92.2%). Performing the time-shift analysis used above on the non-BVCs with significant spatial tuning indicated that most spatially tuned cells of the subiculum code for future positions: Maximum s.i was obtained with reliably positive shifts (Median = 100 ms, *W_sr_* = 35695*, z* = 10.8*, p <* 1 × 10^−27^) as was the maximum ZLAC score (Median = 140 ms, *W_sr_* = 38542*, z* = 12.5*, p <* 1×10^−36^). The magnitude of the time shifts computed from s.i. was marginally larger for BVCs compared to non-BVCs (Median = 120 ms vs. 100 ms, *W_rs_* = 66682*, z* = 1.98*, p* = 0.047) but those computed from ZLAC were not statistically distinguishable (Median = 140 ms vs. 140 ms, *W_rs_* = 66229*, z* = 1.26*, p* = *n.s.*).

### 3.2 BVC Spiking is Better Predicted by Future Position

The above analyses optimize ratemap spatial tuning but do not overtly test that a resulting ratemap better reflects BVC firing. Thus, we also pursued a complementary approach by asking, beyond having greater spatial tuning, does a time-shifted ratemap better predict the

First, to establish a baseline predictability of the BVC spiking, ratemaps were built with no temporal lag or integration of position and we measured the likelihood of the observed spiking when predicted from this ratemap. We then used gradient descent to select a time-lag parameter to optimize the predictive quality of the ratemap as measured by negative log-likelihood. We tested the stability of those estimates with an intra-class correlation analysis and found the parameter estimates were reliable (i.e., broadly consistent) across the k-fold cross-validation runs and between separate recordings (see Table 1). Finally, we compared the different model variants to determine whether the spiking became significantly more predictable given the addition of the time shift to the ratemap generation procedure.

**Table 1:** Intra-class correlations (ICC) for the time lag and time integration parameter estimates across model types to assess fit reliability. Reliability across k-folds are shown in rows labeled with ‘All estimates.’ Reliability across recordings was computed by running ICC on a single cross-validation estimate across recordings for each cell and is shown in rows labeled with ‘Across recs.’ The symbol øis shown for models wherein the specific parameter was not estimated. Abbreviations: dF = degrees of freedom; ICC = Intra-class correlation; int. = integration; recs. = recordings

| ICC values | Integration type: | *None* | *Boxcar* | *Normal* | *Exponential* |
| --- | --- | --- | --- | --- | --- |
|  | *Parameter* | wo / w shift | wo / w shift | wo / w shift | wo / w shift |
| All estimates | Time lag | ∅ / 0.97 | ∅ / 0.96 | ∅ / 0.87 | ∅ / 0.96 |
| | Time int. | ∅ / ∅ | 0.99 / 0.99 | 0.97 / 0.95 | 0.97 / 0.96 |
| Across recs. | Time lag | ∅ / 0.82 | ∅ / 0.88 | ∅ / 0.92 | ∅ / 0.91 |
| | Time int. | ∅ / ∅ | 0.93 / 0.93 | 0.87 / 0.83 | 0.87 / 0.87 |

To compare across models to identify which was best able to predict the BVC spiking activity, we computed the Bayesian Information Criteria (BIC) for each. This standardly used measure rewards good predictions but also applies a penalty for models that do so by using more free parameters, with the final result of selecting the most parsimonious models that perform the best. Lower BIC values are better.

Allowing a temporal shift parameter to be optimized for each ratemap improved the predictive quality of the ratemap. The BIC was significantly reduced (improved) (glme est. [C.I.] = −104 [−119 − 88], *p <* 1 × 10^−38^; see Table 2). The estimated time shifts were reliably positive (Median = 94 ms, *W_sr_* = 1407259*, z* = 19.4*, p <* 1 × 10^−83^; Fig. 4A) indicating that building the ratemap from future position improved how well the ratemap predicted the moment by moment spiking of the BVCs. This is in direct alignment with the results described in the previous section, showing that ratemaps had greater spatial tuning when constructed from future positions.

**Figure 4:**
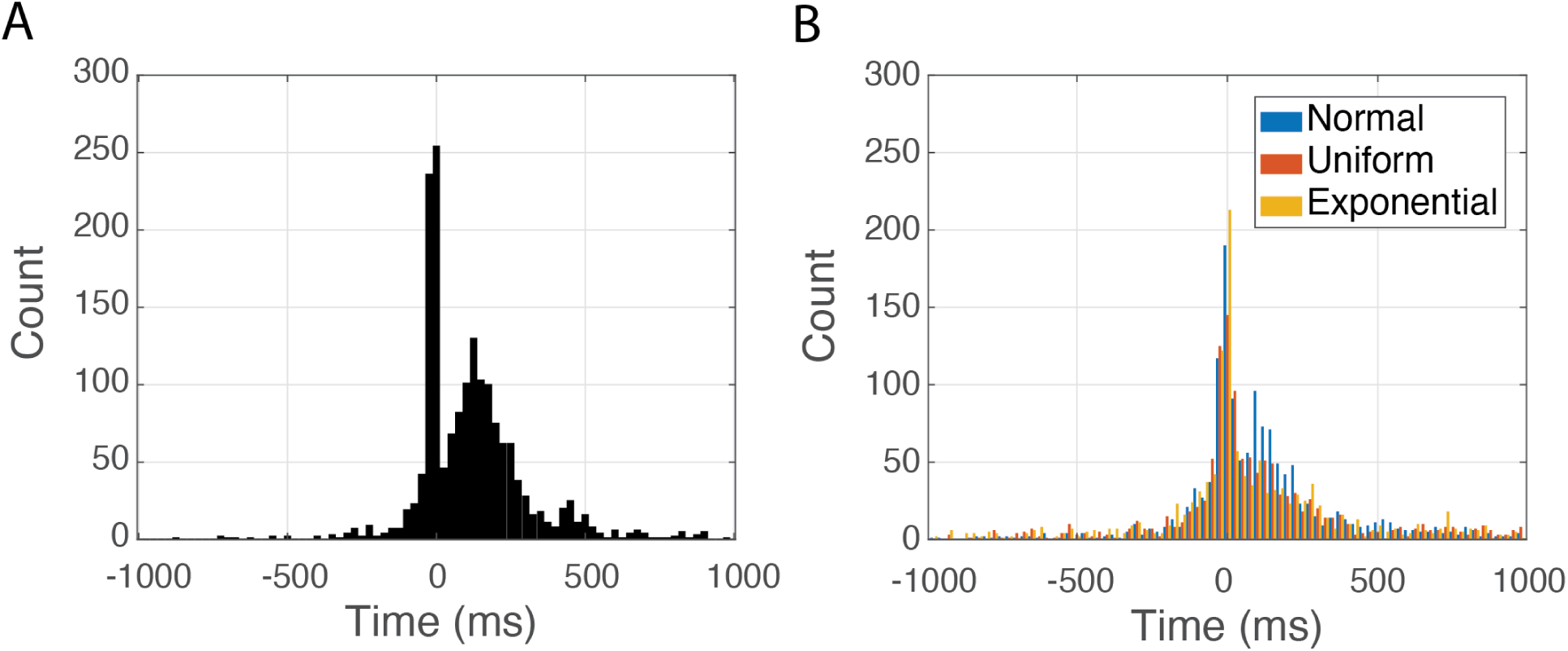
Temporal lags that create ratemaps with the maximum likelihood of generating the observed spiking are reliably positive. A) The histogram of time shifts when no temporal integration is allowed. B) Histograms of time shifts across the three variants of temporal integration. Blue for normally distributed moving average. Orange for uniformly distributed moving average. Yellow for exponentially weighted moving average. Both plots are truncated at 1000 ms for visibility. About 5% of fits were scattered roughly uniformly outside of this window in either direction.

**Table 2:** Summary of maximum likelihood results across model types. BIC is the Bayesian Information Criteria of the standard approach to building ratemaps. Values under ‘effect’ reflect the general linear mixed effect (glme) model estimates of how each modification to how the ratemap is constructed changes the BIC. Negative values indicate improvements in how likely the empirical spiking dynamics were based on the resulting ratemap. Abbreviations: dF = degrees of freedom; C.I. = confidence interval; int. = integration

|  | BIC | effect | dF | C.I. | *p* |
| --- | --- | --- | --- | --- | --- |
| Standard ratemap | 13647 |  |  | [12702 14593] |  |
| + time lag | | -332.79 | 15435 | [-377.33 - 288.25] | $< 10^{-47}$ |
| + normal int. | | -804.66 | 15435 | [-867.65 - 741.67] | $< 10^{-134}$ |
| + boxcar int. | | -890.74 | 15435 | [-953.73 - 827.75] | $< 10^{-164}$ |
| + exponential int. | | -1204.9 | 15435 | [-1267.9 - 1141.9] | $< 10^{-293}$ |

### 3.3 Time-averaged Position Predicts BVC Spiking Better than Instantaneous Position

We next tested whether constructing the ratemaps based on a moving average (i.e., a time-integrated variant) of the position rather than the standard instantaneous position further improved the ability to predict neural spiking. We compared three variants of how the moving average was computes, each differing in how each time point was weighted. A standard moving average weights each point within some window equally and is referred to here as a boxcar weighting. We also used a normally distributed weighting wherein points near an origin are weighted more than those further in the past or future. Finally, an exponentially decaying weighting allowed a bias into either the future or the past. In each, the integration window size was a free parameter, optimized to maximize the likelihood of the spiking based on the ratemap. This was repeated with and without allowing the time shift to also be optimized. As before, we established that the estimates were reliable across cross-validations and recordings (Table 1).

We found that all three forms of temporal integration significantly improved the BIC score (Table 2). The magnitude of the improvement varied across the approaches. The exponentially weighted integration window yielded the largest improvement (Δ*BIC* : exp. vs. no integration = −1204.9 [−1267.9 − 1141.9], *p <* 1 × 10^−293^; see Table 2 for other comparisons). It was significantly better than the other forms of integration (Δ*BIC* : exponential. vs. normally distributed moving average = −400.24 [−463.23 − 337.25], *p <* 1 × 10^−34^; exp. vs. Boxcar moving average = −314.17 [−377.16 − 251.18], *p <* 1 × 10^−21^; not shown in Table 2).

Across all variants of the time-integration approach, the median optimized temporal lags were positive (normal weighting: median = 89 ms, *W_sr_* = 1249300*, z* = 13.0*, p <* 1 × 10^−37^; uniform weighting: median = 43 ms, *W_sr_*= 1111500*, z* = 7.5*, p <* 1 × 10^−13^; exponential weighting: median = 9.2 ms, *W_sr_*= 947450*, z* = 3.9*, p <* 1 × 10^−4^). Distributions of lags over fitted datasets are shown in Figure 4B. In all cases, the distributions are positively shifted but also show a local maximum around zero (i.e., ±20 ms), visible in Figure 4. In the variant without temporal integration (Fig. 4A), the optimized lag for ∼ 23% of datasets was within one frame of zero, ∼ 16% were shifted into the past and ∼ 61% were shifted into the future. Variants with temporal integration had similar distributions ([% *<* ±20 ms, % *<* −20 ms, % *>* 20ms] respectively for normal weighting: [∼ 15%, ∼ 27%, ∼ 58%]; for uniform weighting: [∼ 13%, ∼ 33%, ∼ 54%]; for exponential weighting: [∼ 15%, ∼ 36%, ∼ 48%]).

The distributions of time integration kernel sizes are shown in Figure 5. Across each of the kernel types, a range of values were selected. Across all three kernel types, most of the values align with small magnitudes, indicating that the most predictive ratemaps were generated by limited time integration. However, in each of the cases there was also considerable spread, including values that reflect integration over large temporal windwows. To understand the window of behavior that the optimized parameters align the spiking with, it is important to reconstruct the actual time-integration kernels. The reconstructed time-integration kernels are shown for all optimized parameter sets in Figure 6. across the right-most three columns of panels, a wide variety of kernel widths (marked in orange) can be seen, indicating a spectrum of positional encoding timescales.

**Figure 5:**
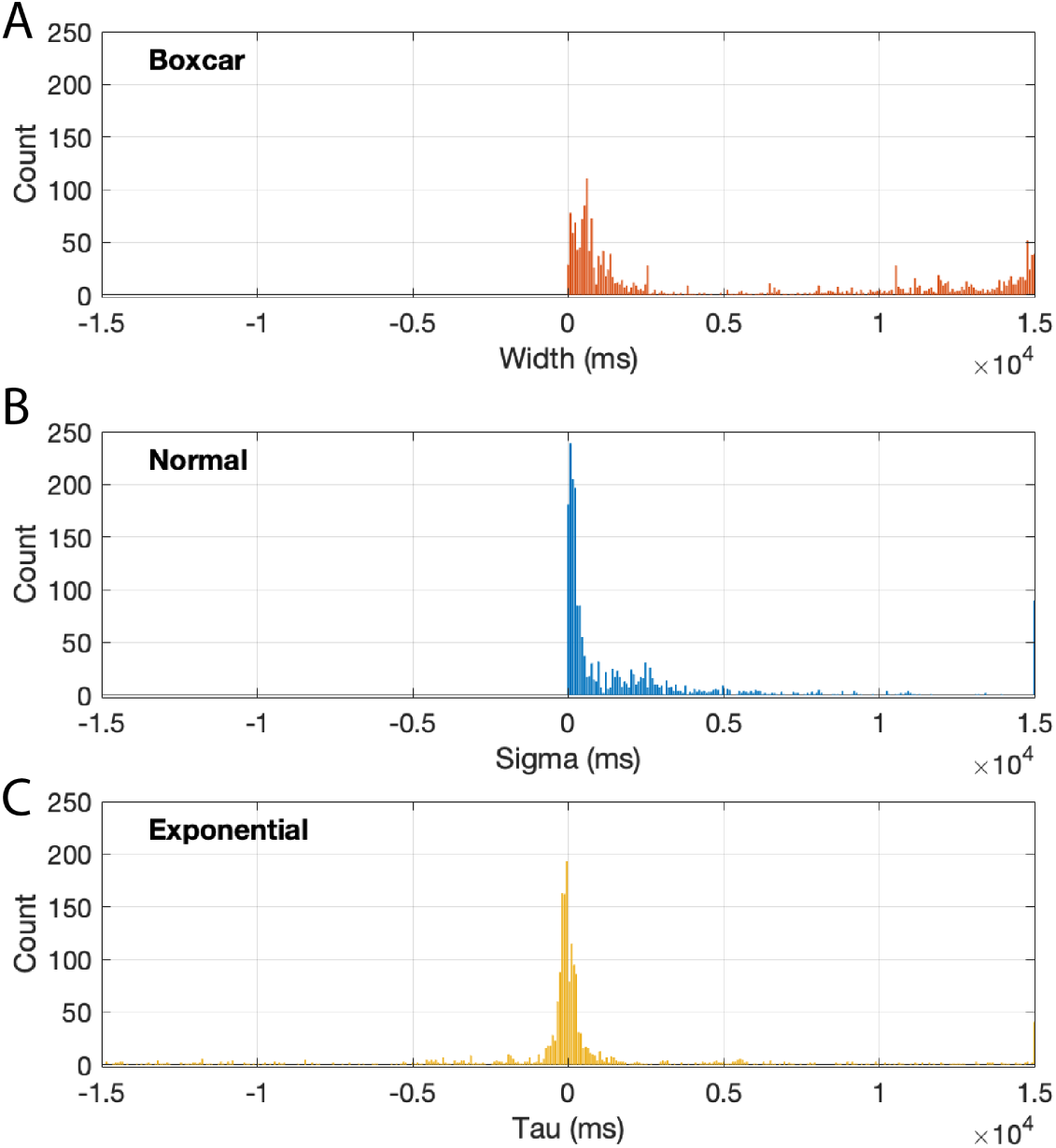
Spectrum of time integration parameters estimates. A) Boxcar widths used in constructing uniformly distributed integration kernels. B) Sigma values used for constructing normally distributed temporal integration kernels. C) Tau equivalents of the omega parameter used to construct the exponentially decaying integration kernel. Positive values constructed kernels that decayed into the future, while negative values constructed kernels that decayed into the past. Note: Values for boxcar width (A) and sigma (B) were constrained to be positive.

**Figure 6:**
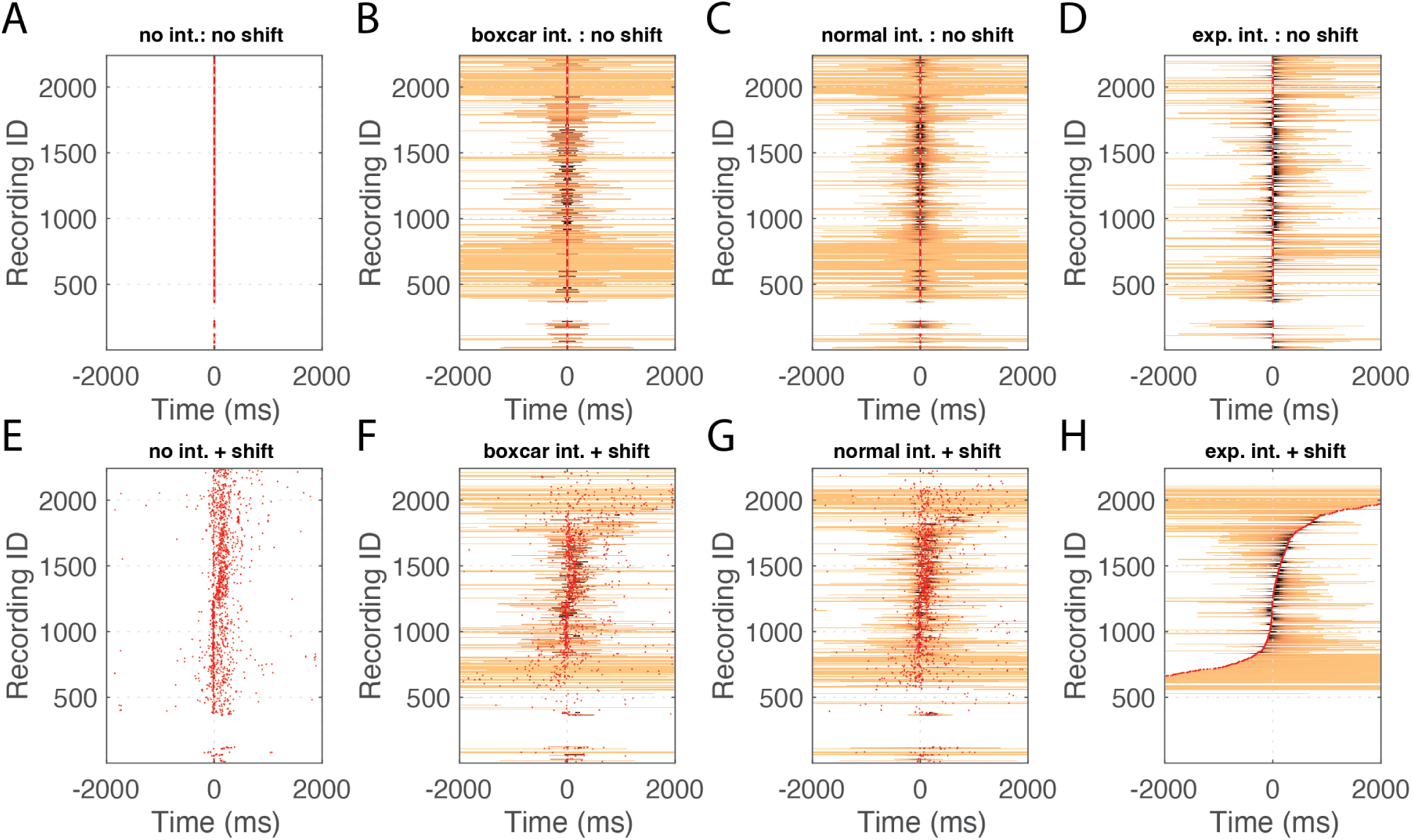
Reconstructed time-integration kernels for all fits across variants. A) With no temporal shift and no integration, all fits were sensitive to the position only at the instant at which a cell fired, reflected by a red dot at zero. B) Reconstructed time-integration kernels for normally distributed weighted moving averages when the temporal shift was fixed at zero. The orange band indicates the width of the kernel. C) Same as B, but for a uniformly distributed weighted moving average. D) Same as B, but for an exponentially distributed weighted moving average. E-H) Same as A-D but for fits for when the temporal lag was optimized as a free parameter. In all plots, the center of the temporal kernel is marked with a red dot for visibility. The x-axes of all plots are truncated at 2000 ms for visibility. About 5% of fits were scattered roughly uniformly outside of this window in either direction. The order of fits is fixed across all panels and is based on a sorting of the exponentially weighted moving average with optimized temporal lag, shown in H.

The reconstructed kernels also illustrate the positive bias of the optimized temporal shifts. Summing over all of the kernels reveals insights into the net sensitivity to position, past and future, across cells. This is shown in Figure 7. Despite the different forms of the time-integration windows used across variants, summing over all kernels of a given type to find the the net sensitivity profile reveals a highly consistent pattern across versions without temporal shifts (Fig. 7A-D) and across versions with temporal shifts (Fig. 7E-H).

**Figure 7:**
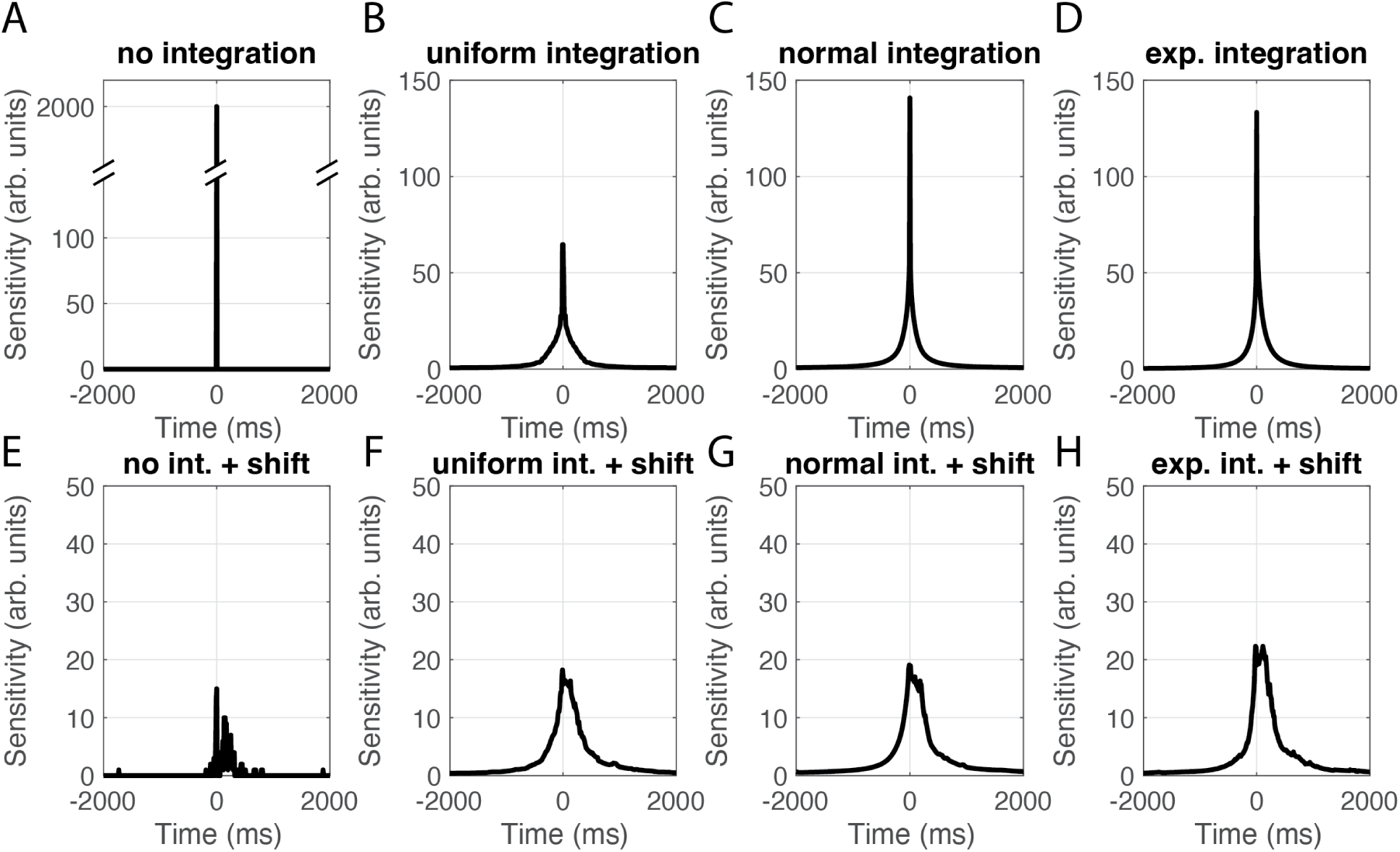
Cumulative reconstructed time-integration kernels across approaches illustrate a consistent sensitivity as a function of time over variants. A) With no temporal shift and no integration, all fits were sensitive to the position only at the instant at which a cell fired, reflected by a peak at zero. B-H) Net sensitivity, computed as the sum over kernels shown in Fig. 6B-H, respectively. Note the changes in the y-axis scale between A, B-D, and E-H.

As a side note, visual inspection of the temporal lags across temporal integration variants (Fig. 4) suggests a trend wherein the percentage of fits with shifts into the past is greater for the exponential integration approach, and the median lag shrinks. However, this is an illusion generated by how the time integration kernels function in practice. The integration windows extend the sensitivity around the instant specified by the time-lag. The exponential kernel, in particular, extends in only one direction from the origin specidied by the time-shift parameter. The net sensitivity profiles (Fig. 7) reveal the interplay between time-shift and the integration window. The resulting net sensitivity remains substantially positively biased. This is seen most easily by re-plotting the same data overlapping shown in Figure 8. Thus, the shifts in temporal lags observed across temporal integration approaches are a reflection of the changing structure of the kernel itself and not inconsistencies across approaches with regard to the optimal alignment between the spiking and behavior. Across the highest performing maximum likelihood based approaches, BVC spiking profiles are reliably best aligned with future position.

**Figure 8:**
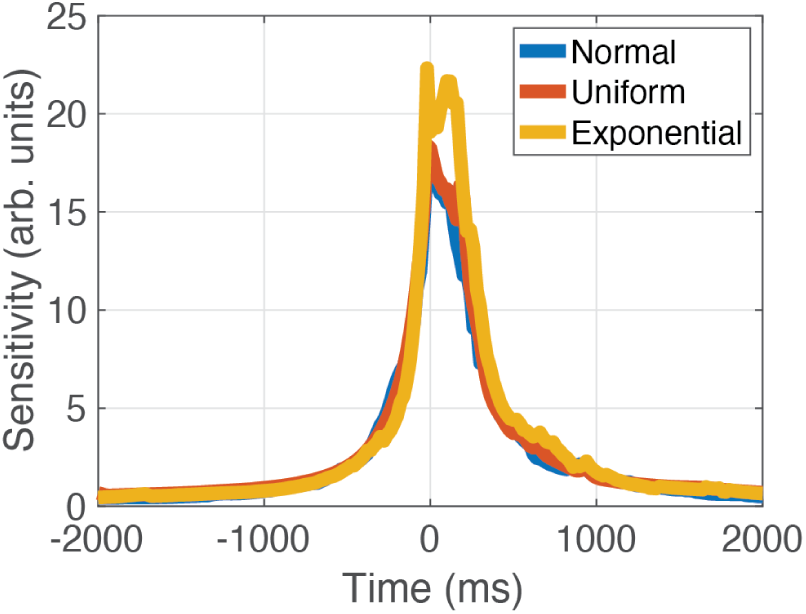
Direct comparison of net sensitivity of time-integration performed across approaches with optimized time shift shows similarities. The lines are replotted from Fig. 7F-H for direct comparability.

### 3.4 Time Shift and Integration Windows of BVCs and Non-BVCs are Similar

Finally, we asked if these tuning properties of BVCs were distinct from non-BVCs. The qualitative pattern of results was overall the same for non-BVCs. Precise statistics can be found in Table 3. The spiking of non-BVCs was significantly more predictable given knowledge of the rat position if a time-shift was allowed. The median parameterized shift was not significantly different from that observed for BVCs (BVC median = 93.9 ms vs. non-BVC median = 98.1 ms, *W_rs_*= 5379183*, z* = 0.40*, p* = 0.69). Following the same trend as with BVCs, the optimized time-shifts remain positive but trend toward shorter magnitudes when temporal integration methods sample future behavior beyond the time shift (Normal median = = 98.1 ms, *W_sr_*= 3751300*, z* = 7.86,, *p <* 1 × 10^−14^; uniform median = 67.3 ms, *W_sr_* = 3544400*, z* = 5.18,, *p <* 1 × 10^−6^) with the exponential integration approach becoming non-significantly above zero (median = 17.9 ms, *W_sr_* = 3334500*, z* = 0.92,, *p* = 0.36). In each case, the optimized temporal shift was not significantly different from that estimated from the BVCs (Normal: BVC median = 88.7 ms vs. non-BVC median = 85.5 ms, *W_rs_*= 5344848*, z* = 0.04*, p* = 0.97; uniform: BVC median = 43.2 ms vs. non-BVC median = 67.3 ms, *W_rs_* = 5276507*, z* = 0.91*, p* = 0.36; exponential: BVC median = 9.2 ms vs. non-BVC median = 17.9 ms, *W_rs_*= 5276507*, z* = 0.91*, p* = 0.36). The optimized time-shifts when there was no temporal integration was allowed were reliably positive (Median = 102.1 ms, *W_sr_*= 3573123*, z* = 14.4*, p <* 1 × 10^−46^). Allowing time-averaging of the position likewise improved the prediction of spiking as shown in Table 3.

**Table 3:** Summary of maximum likelihood results across model types. BIC is the Bayesian Information Criteria of the standard approach to building ratemaps. Values under ‘effect’ reflect the general linear mixed effect (glme) model estimates of how each modification to how the ratemap is constructed changes the BIC. Negative values indicate improvements in how likely the empirical spiking dynamics were based on the resulting ratemap. Abbreviations: dF = degrees of freedom; C.I. = confidence interval; int. = integration

| | BIC | effect | dF | C.I. | $p$ |
| --- | --- | --- | --- | --- | --- |
| Standard ratemap | 13682 |  |  | [12774 14590] |  |
| + time lag | | -179.23 | 26835 | [-188.69 - 169.76] | $< 10^{-47}$ |
| + normal int. | | -752.18 | 26835 | [-765.49 - 738.86] | $\sim 0$ |
| + boxcar int. | | -862.44 | 26835 | [-875.77 - 849.12] | $\sim 0$ |
| + exponential int. | | -795.55 | 26835 | [-808 - 85 - 782.25] | $\sim 0$ |

## 4 Discussion

Spatial coding in the subiculum is less well understood than that in other entorhinal-hippocampal regions such as CA1 and medial entorhinal cortex. See Place et al. (2025); Lee and Lee (2025), and Lever et al. (2025) for subiculum-centred reviews of hippocampal spatial coding. The present study aimed to better understand the temporal coding properties of boundary vector cells (BVCs) recorded in the rat dorsal subiculum. To this end, we analyzed recordings of BVCs previously described by Lever et al. (2009) using two approaches. In the first, we asked if adding a temporal offset between the rat position and the spiking of a BVC increased the apparent spatial tuning in the firing rate map. Finding that it reliably did across BVCs, we asked if the offset that maximized the rate map spatial tuning was consistent with BVCs coding for future versus past positions. We found a consistent bias towards encoding future positions. This pattern of results was mirrored in the second analysis variant that, rather than optimizing spatial tuning in the firing rate map, optimized how well the firing rate map predicted the BVC spiking. Having established that BVCs encode future state, we then asked whether that encoding is focused on a particular temporal scale or, alternatively, captures behavior at multiple scales. To this end, for each cell, we asked “How much time-integration of the behavioral state is the observed spiking most consistent with?” Across cells, we observed a wide spectrum of time-constants of integration, indicating that BVCs form a multiscale encoding of future states. Finally, we compared the distribution of both offsets and integration rates observed across BVCs with those computed by the same approaches for non-BVC cells recorded on the same tetrodes. The distributions were similar, consistent with multiscale encoding of future states being a general property of the subiculum.

These results are, at a high level, consistent with what has been observed in the reciprocally interconnected entorhinal cortex. Entorhinal grid cells, speed cells, and cells of the lateral entorhinal cortex have all been found to exhibit multiscale encoding of state (Chaudhuri-Vayalambrone et al., 2023; Dannenberg et al., 2019; Bright et al., 2020; Tsao et al., 2018). Yet, there are also important differences. Lateral entorhinal cells and speed cells of the medial entorhinal cortex provide multiscale *retrospective* records, whereas we find that BVCs are *prospective*. Entorhinal grid cells are also prospective, yet their hexagonal code affords different computations. Multiscale encodings among grid cells can, in theory (Howard et al., 2014; Shankar et al., 2016), support inferential reasoning about trajectories between distal targets. Multiscale encoding of distance from boundaries, however, may enable reasoning about the timing of events at a boundary.

What computations might an animal perform at environmental boundaries? One clue is that boundary encounters calibrate medial entorhinal grid cells, acting as an error-correction signal (Hardcastle et al., 2015). Such correction demands a calibrated estimate of distance to the boundary. A multiscale encoding of distance to a boundary would provide the necessary information for calculating this correction factor. Consider, for example, the spectrum of tuning curves shown in Figure 6H. Across a population of cells, there would be graded information about when a boundary was reached. In the moments leading up to and following an interaction with a boundary, there would be sufficient information across the population of BVCs to infer the relative trajectory taken. That estimate could then update the animal’s current positional belief or perceived environmental scale.

The future bias of BVCs aligns them with the prospective coding of other spatially tuned neurons of the hippocampal formation (Muller and Kubie, 1989; Chaudhuri-Vayalambrone et al., 2023). Place cells, grid cells, and BVCs all tend to fire 100 − 200ms before the animal reaches a salient location. This prospective bias is in accord with the hypothesis that the hippocampus encodes a successor representation (Stachenfeld et al., 2017; Momennejad, 2020; Geerts et al., 2023). At its core, the successor-representation view holds that hippocampal activity encodes *what comes next* —not a static map of space or context—conditioned on the current state and habitual transitions. The consistent direction and scale of prospective coding among spatially tuned neurons distributed across the hippocampal formation, including CA1, MEC, and subiculum, speak to the development of the successor representation being a circuit-wide computation.

Multiscale encoding among spatially tuned neurons, including the BVCs examined here, in the context of a successor-representation view of hippocampal function, may align with temporal discounting. Temporal discounting is the idea that future states that will occur after a long delay are encoded more weakly than those that will occur soon but, importantly, are nonetheless encoded. This is, in effect, a way to interpret the multiscale integration windows we observed. Though many BVCs encoded boundaries that will be reached shortly, there were nonetheless BVCs that responded considerably earlier.

Departing from the successor-representation view, however, we also observed substantial retrospective coding. In 15 − 30% of our fits, across approaches, we found that the temporal lags that made the empirical spiking most likely were offset into the past. Assuming that this does not reflect measurement error, this is not well accounted for by a standard successor-representation perspective. Minor modifications could reconcile this mismatch.

The Tolman-Eichenbaum Machine (Whittington et al., 2020), for example, contains elements of the Successor-representation, including having an imposed bias toward predictive coding. Yet, it also includes associative learning that will form bidirectional associations between states. These associations could, in principle, reactivate prior states given a sensory cue, thereby introducing some retrospective coding.

The present work provided consistent evidence of future-biased multiscale encoding of position based on the recordings from Lever et al. (2009), but there are several limitations important to acknowledge. One is that the present work did not overtly model the boundary-oriented nature of the coding itself. Rather, the rate map-based approach is agnostic to the form of spatial coding. Though this adds a level of generality, the generality runs the risk of missing key elements of the temporal tuning that may only appear in analyses optimized for boundary coding. Part of the reason for not taking that approach here is that accurate modeling of interactions with boundaries would require precise records of the timing of the moments of perceiving a boundary, which is far from straightforward to obtain. Finally, it is worth recognizing that there is no ground-truth for the designations of BVC and non-BVC. Consequently, cells may have been mislabeled in either direction. The risk of which, as it relates to the current work, is that temporal shifts observed for non-BVCs may nonetheless reflect BVC coding and that some portion of the heterogeneity in temporal coding observed over BVCs was introduced by inclusion of non-BVCs. Given the careful comparison of spatial coding over multiple sessions per cell performed by Lever et al. (2009), however, we do not expect this risk to be high, particularly with regard to the possible mis-classification of non-BVCs as BVCs.

In conclusion, the current work showed through the use of multiple complementary analyses that boundary vector cells, BVCs, are biased toward encoding future positions and that encoding is most consistent with a time-integrated version of position. Across cells, we observed variability in the width of the window over which that integration was performed. From these findings, we conclude that BVCs encode a future-biased spectrum of positions in the rat.

## Acknowledgements

We gratefully acknowledge support from the National Institutes of Health’s National Institute on Aging, grant R01AG076198 to ZT and ELN. CL gratefully acknowledges grant funding from the BBSRC (BB/T014768/1). This research was also supported in part by Lilly Endowment, Inc., through its support for the Indiana University Pervasive Technology Institute and by Shared University Research grants from IBM, Inc., to Indiana University.

## Conflicts of interest

The authors have no conflicts interests to declare.

## Artificial Intelligence Generated Content (AIGC)

ChatGPT, models 3.5 and 4 by OpenAI, was used to assist in the development and debugging of analyses code and for producing the LaTeX markup for equations and tables appearing in this manuscript.

## Notes

### Competing Interest Statement

The authors have declared no competing interest.

